# Development of a genetically encoded fluorescent indicator for monitoring GPR52 activation

**DOI:** 10.64898/2026.03.11.711017

**Authors:** Guangyi Lan, Huan Wang, Tongrui Qian, Shu Xie, Cheng Qian, Daniel Ursu, Klaus D. Bornemann, Bastian Hengerer, Yulong Li

**Affiliations:** State Key Laboratory of Membrane Biology, Peking University School of Life Sciences, Beijing 100871, China; PKU-IDG/McGovern Institute for Brain Research, Beijing 100871, China; Peking-Tsinghua Center for Life Sciences, New Cornerstone Science Laboratory, Academy for Advanced Interdisciplinary Studies, Peking University, Beijing 100871, China; National Biomedical Imaging Center, Peking University, Beijing 100871, China; Neuroscience & Mental Health, Boehringer Ingelheim Pharma GmbH & Co. KG, Biberach, Germany

**Author notes:** Manuscript correspondence: Yulong Li.

## Abstract

GPR52 is an orphan G protein-coupled receptor implicated in psychiatric and neurodegenerative disorders, but its endogenous ligand remains unidentified, limiting the exploration of its physiological functions and therapeutic potential. We pioneered a novel methodology for orphan GPCR ligand discovery utilizing the GPCR-activation-based (GRAB) strategy by developing GPR52-1.0, a genetically encoded fluorescent sensor. GPR52-1.0 exhibits excellent membrane trafficking and high sensitivity in HEK293T cells, cultured neurons, and acute mouse brain slices. Notably, it detects neuronal activity–dependent endogenous ligand release in the striatum, with responses abolished by a specific antagonist. This sensor provides a powerful tool for identifying GPR52’s endogenous ligand(s) and enables real-time monitoring of its activation. Our work lays the foundation for uncovering GPR52’s physiological roles and supports future efforts to develop GPR52-targeted therapeutics.

## INTRODUCTION

G protein-coupled receptors (GPCRs) represent one of the largest and most versatile families of membrane proteins, characterized by seven-transmembrane domains and activation by diverse extracellular ligands^1^. Upon ligand binding, GPCRs initiate intracellular signaling cascades that regulate a wide array of processes across organ systems. As such, GPCRs constitute the largest class of drug targets, with over one-third of FDA-approved drugs acting on this receptor family^2^. In the nervous system, GPCRs are abundantly expressed and critically involved in modulating synaptic transmission, neural excitability, and behavior outputs^3^. These features make them particularly important for understanding brain function and treating neurological and psychiatric conditions.

Despite substantial progress in GPCR research, a significant number of GPCRs remain classified as orphan receptors, with their endogenous ligands unknown^4^. This knowledge gap limits the mechanistic understanding of these receptors and hinders their therapeutic exploitation. Among these orphan GPCRs, GPR52 has emerged as a particularly interesting candidate due to its involvement in neuropsychiatric disorders—including schizophrenia and anxiety—and its potential neuroprotective roles in Huntington’s disease, as shown in both genetic and pharmacological studies^5-8^.

Notably, GPR52 differs from many classical ligand-dependent GPCRs in exhibiting a high level of constitutive G/cAMP signaling in the absence of an exogenous agonist^9^. Previous functional studies demonstrated substantial basal signaling by GPR52, and subsequent structural analyses revealed that extracellular loop 2 (ECL2) occupies the canonical orthosteric pocket and functions as a built-in agonist, thereby stabilizing a self-activated receptor conformation^10^. Nevertheless, GPR52 can be further modulated by synthetic ligands that bind to a distinct extracellular side pocket^10^, indicating that its constitutive activity does not preclude additional dynamic regulation. Whether endogenous molecules dynamically modulate GPR52 in native tissues remains unknown, which has posed a major obstacle to further functional and translational exploration.

To address this challenge, we employed the GPCR-Activation-Based (GRAB) strategy^11,12^ to develop a genetically encoded fluorescent sensor based on GPR52. In this study, we report the engineering, optimization, and validation of the GPR52-1.0 sensor in live cells and intact brain tissues. We demonstrate its utility in detecting synthetic ligands and, importantly, the sensor revealed endogenous ligand release upon electrical stimulation in the striatum. These findings establish GPR52-1.0 as a tool for investigating the dynamic regulation of GPR52 and provide a basis for future identification and characterization of endogenous GPR52 modulators, while also opening new avenues for studying its physiological functions and facilitating GPR52-targeted drug discovery.

## RESULTS

### Development of a genetically encoded GPR52 sensor

To enable real-time monitoring of GPR52 modulation, we leveraged the GRAB strategy to develop a genetically encoded fluorescent sensor capable of reporting GPR52 activation (**Fig. 1A**)^13^. Specifically, we first replaced the ICL3 in the human GPR52 receptor with the ICL3 in already existing sensor GRAB_NE1m_ to generate a prototype green GPR52 sensor, and then systematically optimized the linker sequences flanking the circularly permutated green fluorescent protein (cpEGFP) and key residues influencing fluorescence intensity and protein folding in the cpEGFP (**Fig. 1B**). Through the screening of ∼800 sensor variants, we identified GPR52-1.0 as the top-performing construct, which exhibits the highest response (ΔF/F0) to the application of a selective GRP52 agonist^14^ (**Fig. 1B**).

**Fig. 1.**
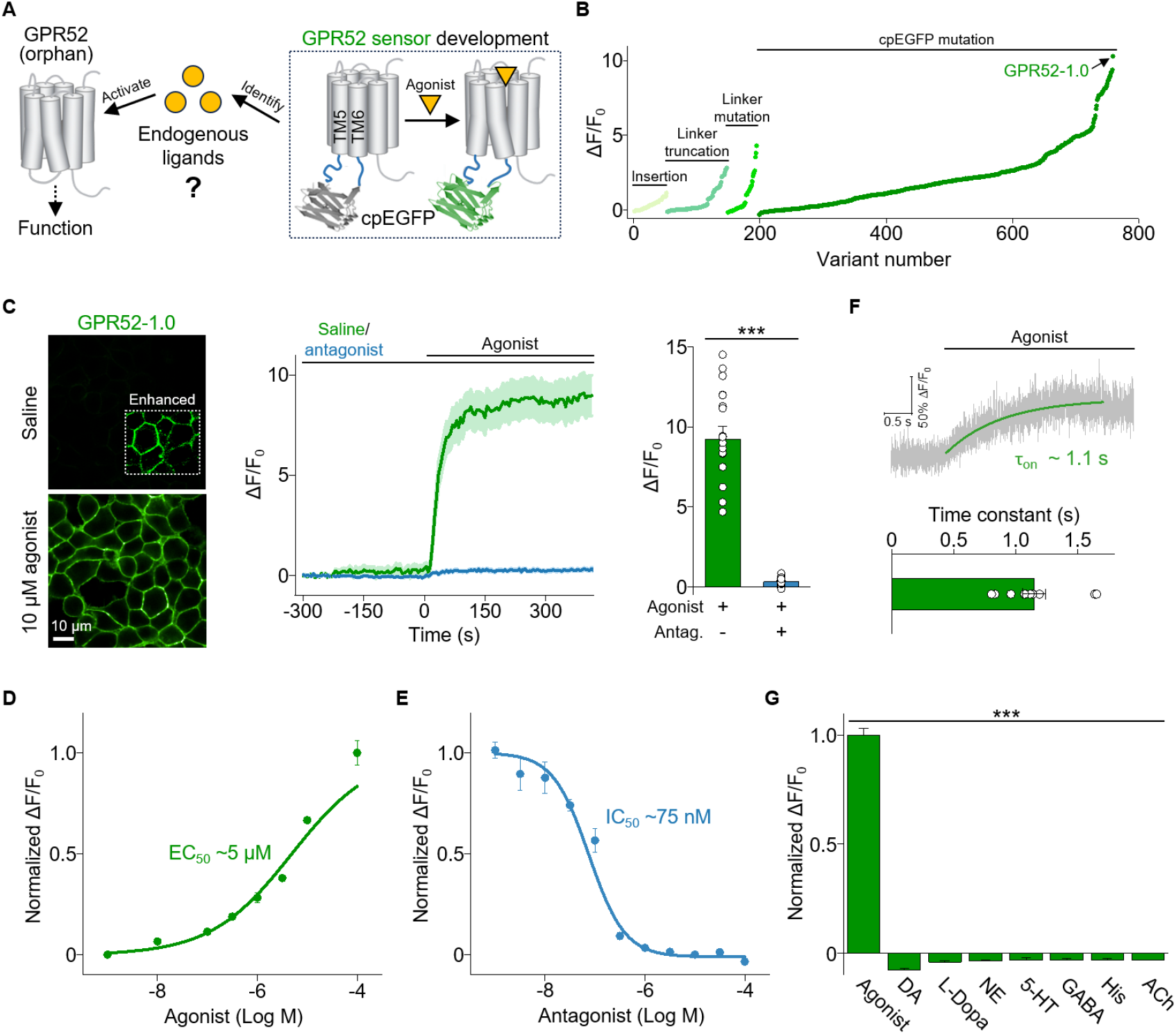
Development and characterization of GPR52 sensors in HEK293T cells. (**A**) Schematic drawing shows the strategy of developing GRAB GPR52 sensors to identify endogenous ligands of GPR52. (**B**) Screening and optimization steps of GRAB GPR52 sensors and the fluorescence response to 10 μM synthetic agonist; the black arrow indicates the best candidate GPR52-1.0. (**C**) Example images (left), traces (middle) and quantification (right) of the change in GPR52-1.0 fluorescence in response to 10 μM agonist with (blue) or without (green) antagonist pretreatment; n = 17–20 cells from 3 coverslips; ^***^*p* < 0.001; Two-Sample *t*-test. (**D**) Normalized dose–response curves measured in HEK293T cells expressing GPR52-1.0, with the corresponding EC_50_ value for agonist shown; n = 3 repeats. (**E**) Normalized dose–response curves measured in HEK293T cells expressing GPR52-1.0, with the corresponding IC_50_ value for antagonist in the presence of 10 μM agonist shown; n = 3 repeats. (**F**) Top, representative fluorescence changes in GPR52-1.0-expressing cells in response to the local perfusion (100 μM agonist in pipette with normal bath solution). Bottom, group data summarizes on time constants measured upon application of agonist; n = 10 cells from 3 coverslips. (**G**) Normalized fluorescence change in response to the indicated compounds (each at 10 μM) measured in cells expressing GPR52-1.0. DA, dopamine; L-Dopa, levodopa; NE, norepinephrine; 5-HT, 5-hydroxytryptamine; GABA, γ-aminobutyric acid; His, histamine; ACh, acetylcholine; n = 3 repeats; ^***^*p* < 0.001; One-Way ANOVA. In this and subsequent figures, unless indicated otherwise summary data are presented as the mean±SEM.

When expressed in HEK293T cells, GPR52-1.0 was efficiently localized to the plasma membrane and exhibited robust fluorescence increases in response to the GPR52 agonist. This fluorescence increase was almost completely abolished upon co-application of a BI-derived GPR52 inverse agonist^15^ (**Fig. 1C**). Dose–response analysis revealed an apparent half-maximal effective concentration (EC_50_) of 5 μM for the agonist and a half-maximal inhibitory concentration (IC_50_) of 75 nM for the antagonist (**Fig. 1D and 1E**). High-speed line scan imaging demonstrated rapid fluorescence increases upon local puffing of agonist, with an average activation time constant of ∼1.1 s (**Fig. 1F**). Notably, this sensor showed high specificity, showing no response to a panel of neurotransmitters (**Fig. 1G**). Collectively, these results indicate that GPR52-1.0 is a sensitive and selective tool for monitoring GPR52 activation in living cells.

### Characterization of GPR52-1.0 in cultured neurons and acute mouse brain slices

We next expressed GPR52-1.0 in cultured rat cortical neurons and acute mouse brain slices for further characterization. In cultured neurons, GPR52-1.0 exhibited efficient membrane trafficking and retained fluorescence responses, ligand affinity, and specificity comparable to those observed in HEK293T cells (**Fig. 2A–2C**). To evaluate sensor performance in intact brain tissue, we delivered GPR52-1.0 to the mouse striatum—a brain region with high endogenous GPR52 expression^5^—via adeno-associated virus (AAV) (**Fig. 2D**). In acute brain slices containing striatum, we observed robust fluorescence signals in response to agonist perfusion, which were blocked by antagonist treatment (**Fig. 2D**). Collectively, these data suggest that GPR52-1.0 is functional in native neuronal environments and suitable for detecting ligand-induced activation in brain tissue.

**Fig. 2.**
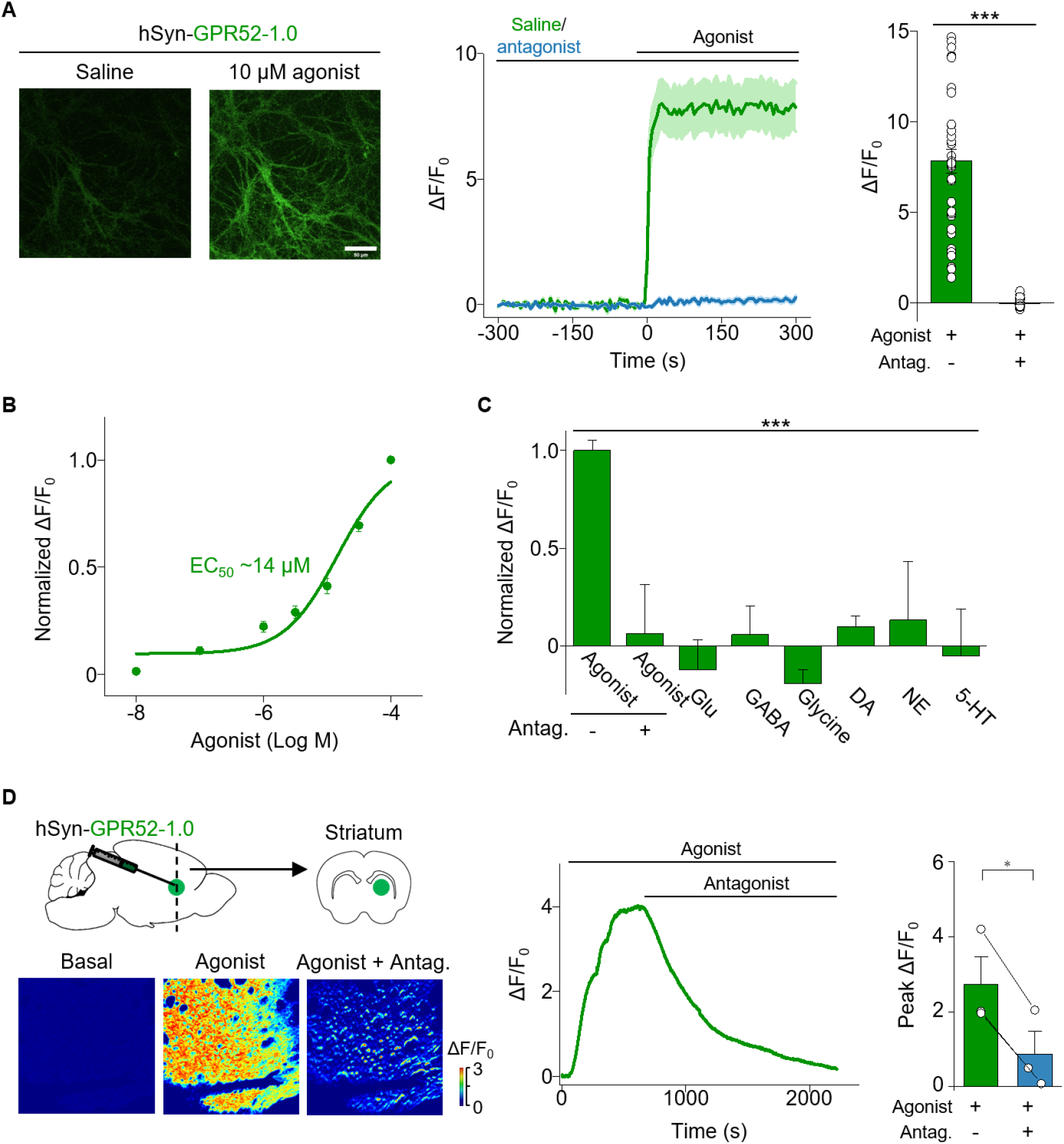
Characterization of GPR52-1.0 in cultured neurons and acute mouse brain slices. (**A**) In GPR52-1.0-expressing neurons, example images (left), traces (middle) and quantification (right) of the change in GPR52-1.0 fluorescence in response to 10 μM agonist with (blue) or without (green) antagonist pretreatment; n = 35–36 neurons from 3 coverslips; ^***^*p* < 0.001; Two-Sample *t*-test. (**B**) Normalized dose–response curves measured in neurons expressing GPR52-1.0, with the corresponding EC_50_ value for agonist shown; n = 3 repeats. (**C**) Normalized fluorescence change in response to the indicated compounds (each at 10 μM) measured in neurons expressing GPR52-1.0. When indicated, the antagonist was also added. Glu, glutamate; n = 3 repeats; ^***^*p* < 0.001; One-Way ANOVA. (**D**) Left: top, schematic illustration depicting the experimental design of virally expressing GPR52-1.0 in the striatum; bottom, representative pseudo-color images of the fluorescence change in GPR52-1.0-expressing acute mouse brain slices; where indicated, 10 μM agonist and 10 μM antagonist were applied. Middle, a representative trace; where indicated, agonist and antagonist were added. Right, summary data; n = 3 slices from 3mice; s; ^*^*p* < 0.05; Paired *t*-test.

### GPR52-1.0 reports endogenous GPR52 ligand release

To investigate whether GPR52-1.0 can detect endogenously released ligands, we virally expressed the sensor in the mouse striatum and prepared acute brain slices 3 weeks post-injection (**Fig. 3A**). Upon electrical stimulation, we observed robust fluorescence increases of GPR52-1.0 (**Fig. 3B1–3B3**). Critically, these responses were significantly reduced by co-application of the GPR52 antagonist, indicating that the observed signal was specific to GPR52 activation (**Fig. 3B2–3B3**). Taken together, these results support the existence of neuronal activity-dependent endogenous molecules capable of activating GPR52 in the striatum.

**Fig. 3.**
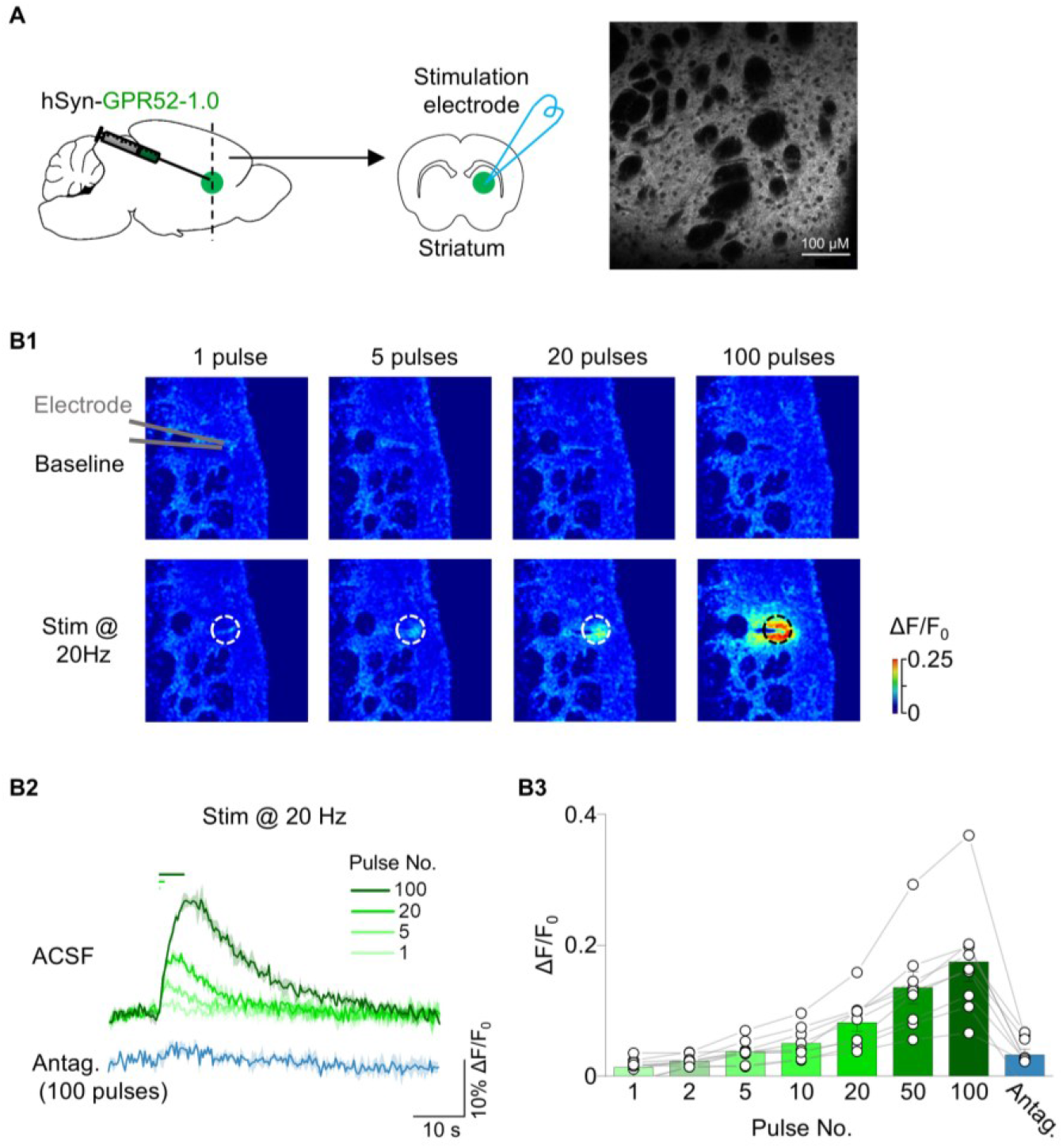
GPR52-1.0 can report the release of GPR52 endogenous ligands *ex vivo*. (**A**) Left, schematic illustration depicting the experimental design in the striatum for (B1–B3); right, a representative image of GPR52-1.0-expressing brain slices. (**B1**) Representative pseudo-color images of GPR52-1.0-expressing brain slices at baseline (top) and in response to the indicated stimuli (bottom) in the presence of artificial cerebrospinal fluid (ACSF). The dashed circles indicate the ROI used to calculate the response, and the approximate location of the stimulating electrode is indicated. (**B2–B3**) Representative traces (B2) and summary data (B3) for the change in GPR52-1.0 fluorescence in response to the indicated stimuli in ACSF or antagonist; n = 9 slices from 3 mice.

## DISCUSSION

Orphan GPCRs, including GPR52, remain underexplored despite their therapeutic relevance in brain disorders. Identifying endogenous ligands is essential not only for understanding their native signaling mechanisms but also for unlocking new drug targets. In this study, we report the development of a genetically encoded fluorescent sensor, GPR52-1.0, which enables dynamic visualization of GPR52 activation in living cells and intact brain tissue. Critically, we pioneered a novel methodology for orphan GPCR ligand discovery utilizing GRAB sensors. The sensor revealed endogenous ligand-induced GPR52 activation upon electrical stimulation in mouse striatal slices.

By enabling real-time tracking of GPR52 activation, GPR52-1.0 serves as a valuable tool to guide biochemical purification or genetic screens for endogenous ligands. Such discoveries could advance our understanding of GPR52’s role in brain function and its contribution to neuropsychiatric and neurodegenerative diseases. Furthermore, the sensor offers a high-throughput platform for screening pharmacological modulators, accelerating the development of GPR52-targeted therapies. More broadly, this approach exemplifies how GRAB-based sensors can facilitate the deorphanization of GPCRs, advancing both basic neuroscience and translational drug discovery.

Conventional pharmacological characterization of GPR52 has primarily relied on downstream cAMP-based assays, including HTRF and GloSensor measurements^7,8^. While these approaches provide sensitive and scalable readouts of G signaling, GPR52-1.0 offers a complementary, receptor-proximal readout by converting receptor conformational changes directly into fluorescence changes without relying on downstream second-messenger accumulation. This distinction is particularly relevant for GPR52, which exhibits high constitutive Gs/cAMP activity that can contribute substantially to the basal signal in downstream assays^16^. In contrast, GPR52-1.0 was optimized based on agonist-evoked ΔF/F, rather than basal fluorescence or basal signaling, and is therefore designed to resolve dynamic modulator-induced changes relative to its own baseline. Together with its second-scale kinetics and spatially resolved optical readout, these properties make GPR52-1.0 particularly suitable for monitoring dynamic GPR52 modulation *in vivo* and *in vitro*.

While our current study stops short of fully deorphanizing GPR52, the sensor we developed lays the groundwork for achieving this in future experiments. The observed activity-dependent activation in striatal slices strongly suggests the presence of endogenous ligands, and GPR52-1.0 provides a robust platform to identify them. A logical next step would involve biochemical fractionation of brain tissue extracts, particularly from the striatum, followed by activity-guided screening using the sensor to detect fractions that elicit fluorescence responses. These active fractions could then be analyzed using mass spectrometry to identify candidate molecules. Together, these strategies would enable systematic identification and validation of endogenous GPR52 ligands, ultimately advancing the deorphanization process and opening new avenues for therapeutic exploration.

## METHODS

### Animals

C57BL/6N mice (6 to 8 weeks of age) were obtained from Beijing Vital River Laboratory Animal Technology Co., Ltd., and group-housed under a 12-hours/12-hours light/dark cycle with a 25°C ambient temperature. All animal experiments were approved by the Animal Care and Use Committee of Peking University School of Life Sciences.

### Compounds

The GPR52 agonist used here was synthesized based on the structure described in WO 012020738^14^. The GPR52 antagonist was previously published^15^. Both compounds were synthesized and provided by Boehringer Ingelheim. Compounds were dissolved in water or DMSO at stock concentrations of 10 mM or 100 mM and stored at -20 °C.

### Cell culture

HEK293T cells (CRL-3216, ATCC) were used to express and test sensors. All cell lines were cultured in DMEM (Gibco) supplemented with 10% (vol/vol) FBS (Gibco) and 1% penicillin-streptomycin (Gibco) at 37 °C in 5% CO2.

### Primary neuronal cultures

Rat cortical neurons were cultured from postnatal day (P) 0 Sprague–Dawley rat pups of both sexes (Beijing Vital River). Specifically, the brain was removed and the cortex was dissected; neurons were then dissociated in 0.25% trypsin-EDTA (Gibco), plated on 12-mm glass coverslips coated with poly(d-lysine) (Sigma-Aldrich) and cultured in neurobasal medium (Gibco) containing 2% B-27 supplement, 1% GlutaMax (Gibco) and 1% penicillin-streptomycin (Gibco) at 37 °C in 5% CO_2_.

### Fluorescence imaging of cultured cells

An inverted confocal microscope (Nikon) equipped with NIS-Elements 4.51.00 software (Nikon), a ×40/1.35-NA oil-immersion objective, a 488-nm laser and a 561-nm laser was used for imaging; the GFP and RFP signals were collected using 525/50-nm and 595/50-nm emission filters, respectively. Cultured cells expressing GRAB GPR52 sensors were either bathed or perfused with Tyrode’s solution containing (in mM) 150 NaCl, 4 KCl, 2 MgCl_2_, 2 CaCl_2_, 10 HEPES and 10 glucose (pH 7.4); where indicated, drugs and other compounds were delivered via a custom-made perfusion system or via bath application. An Opera Phenix high-content screening system (PerkinElmer) equipped with a ×40/1.1-NA water-immersion objective, a 488-nm laser and a 561-nm laser was also used for imaging; the GFP and RFP signals were collected using 525/50-nm and 600/30-nm emission filters, respectively. The Harmony 4.9 software of the Opera Phenix high-content screening system (PerkinElmer) was used for data collection. For imaging, the fluorescence signals of the candidate GRAB GPR52 sensors were calibrated using the GFP/RFP fluorescence ratio. To measure the response kinetics of the GPR52-1.0 sensor, the line-scanning mode of the confocal microscope was used to record rapid changes in fluorescence; a glass pipette containing 100 µM agonist was placed near the surface of HEK293T cells expressing GPR52-1.0, and agonist was puffed onto cells to measure τ_on_.

### Preparation and fluorescence imaging of mouse acute brain slices

Wild-type C57BL/6N mice were deeply anesthetized by an intraperitoneal injection of avertin (500 mg kg^−1^; Sigma-Aldrich) and then placed in a stereotaxic frame for injection of AAVs using a microsyringe pump (Nanoliter 2000 Injector, WPI). hSyn-GPR52-1.0 AAVs were injected (300 nl) into the striatum of mice using the following coordinates: anteroposterior (AP), +1.0 mm relative to bregma; mediolateral (ML), −1.0 mm; dorsoventral (DV), −3.1 mm from the dura.

Three weeks after viral injection, mice were again deeply anesthetized with an intraperitoneal injection of avertin and transcardial perfusion was performed using cold oxygenated slicing buffer containing (in mM) 110 choline chloride, 2.5 KCl, 1 NaH_2_PO_4_, 25 NaHCO_3_, 7 MgCl_2_, 25 glucose, 0.5 CaCl_2_, 1.3 sodium ascorbate and 0.6 sodium pyruvate. Brains were then rapidly removed and immersed in the oxygenated slicing buffer, after which the cerebellum was trimmed using a razor blade. The brains were then glued to the cutting stage of a VT1200 vibratome (Leica) and sectioned into 300-μm-thick coronal slices. Brain slices containing the striatum were incubated at 34 °C for at least 40 min in oxygen-saturated artificial cerebrospinal fluid (ACSF) buffer containing (in mM) 125 NaCl, 2.5 KCl, 1 NaH_2_PO_4_, 25 NaHCO_3_, 1.3 MgCl_2_, 25 glucose, 2 CaCl_2_, 1.3 sodium ascorbate and 0.6 sodium pyruvate. For two-photon imaging, the slices were transferred into an imaging chamber in an FV1000MPE (Olympus) microscope equipped with a ×25/1.05-NA water-immersion objective and a mode-locked Mai Tai Ti:Sapphire laser (Spectra-Physics) tuned to 920 nm with a 495-to 540-nm filter to measure fluorescence. FV10-ASW Ver.3.1a software for the FV1000MPE two-photon microscope (Olympus) was used for data collection. For electrical stimulation, a homemade bipolar electrode (WE30031.0A3, MicroProbes) was placed onto the surface of the brain slice near the striatum under fluorescence guidance. Imaging and stimulation were synchronized using an Arduino board with a custom-written program. All other stimulation experiments were recorded at video frame rates of 0.3583 s per frame, with 256 × 192 pixels per frame. The stimulation voltage was set at 5–8 V, and the duration of each stimulation was 1 ms. Where applicable, compounds were applied to the imaging chamber by perfusion in ACSF at a flow rate of 4 ml min^−1^.

### Statistical analysis

Statistical analyses were performed using OriginPro 2020. Group data were analyzed using Two-Sample *t*-Test, Pair-Sample *t*-Test, One-Way ANOVA, or Mann-Whitney *U* test, and differences were considered significant at *p* < 0.05. Unless indicated otherwise, all summary data are presented as the mean±SEM.

## Data and code availability

All data supporting the findings of this study are available within the main text or the Supplementary Information.

